# Emotional Mirrors in the Rat’s Anterior Cingulate Cortex

**DOI:** 10.1101/450643

**Authors:** M Carrillo, Y Han, F Migliorati, M Liu, V Gazzola, C Keysers

**Affiliations:** Social Brain Lab, Netherlands Institute for Neuroscience, Royal Netherlands Academy of Arts and Sciences, Meibergdreef 47, 1105BA Amsterdam, the Netherlands; Department of Psychology, University of Amsterdam, Amsterdam, The Netherlands; Current Address: Cognitive Psychology Unit, Institute of Psychology, Leiden University, Leiden, The Netherlands

## Abstract

How do the emotions of others affect us? The human anterior cingulate cortex (ACC) responds while experiencing pain in the self and witnessing pain in others, but underlying cellular mechanisms remain poorly understood. Here we show the ACC (area 24) contains neurons responding when a rat experiences pain and while witnessing another receive footshocks. Most of these do not respond to a fear conditioned tone (CS). Deactivating this region reduces freezing while witnessing footshocks to others but not while hearing the CS. A decoder trained on spike counts while witnessing footshocks can decode the animal’s own pain intensity when experiencing pain. Mirror-like neurons thus exist in ACC that encode the pain of others in a code shared with pain but not fear in the self.

**One Sentence Summary:** ACC contains neurons responding selectively when a rat witnesses another’s pain and experiences pain in the self.

## Introduction

Understanding how we share the affective state of others is important for understanding social interactions [1]. Neuroimaging shows humans recruit their anterior cingulate cortex (ACC) both while experiencing pain and, vicariously, while witnessing pain in others [2]. This vicarious activity is stronger in more empathic individuals [2,3] and reduced in psychopathy [4], making it a candidate for a neural mechanism of affect sharing. Some argue, these neuroimaging findings reflect the existence of pain mirror neurons, i.e. neurons selectively responding during the experience of pain and the perception of other people’s pain [5]. That some ACC neurons respond to the observation and experience of pain is supported by reports of one such neuron in a human patient [6] and by one report of neurons in the mouse ACC in which the early gene *arc* is more expressed following the experience of footshocks and witnessing another animal receive footshocks [7]. The functional properties of these neurons however remain unknown. A pivotal question regards the selectivity of responses in the ACC[8,9]. It has been argued that a vicarious response in the ACC can only signal that someone else is in pain if it has at least the following two features [9]. First, neural responses must be *selective*. If the same neuron responds to the experience of pain and of other salient emotions (e.g. fear), its firing cannot signal pain as different from these other emotions[8,9]. Second, the population of neurons should employ a *common code* to signal pain in the self and in others: if the brain reads out the pain of others from the vicarious ensemble activation of a subset of its own pain neurons, a decoder able to decode pain levels of others from ensemble activity should be able to decode pain levels in the self from the same ensemble using the same rule [10,11]. Unfortunately, fMRI experiments have failed to provide robust evidence for either of these criteria despite intense efforts to do so. Instead, the literature shows that the ACC is recruited by many salient stimuli beyond pain [8,9] and that the pattern of activity signaling pain in others differs from that signaling pain in the self [9,10, but see 11]. That functional neuroimaging pools the activity of millions of neurons within each voxel may cause this failure. Here, we therefore use a rodent model of emotional contagion in which a shock experienced animal witnesses a conspecific experience painful electroshocks [12–18] to explore this issue at this cellular level. We chose rodents as a model because we know that region 24 of the rodent ACC, including regions Cg1 and Cg2, is similar in cytoarchitecture and connectivity to the region implicated in pain empathy in humans[2,9,19] and that the ACC of rodents is activated by the distress of others [7,14,20]. To explore the specificity and common coding of neurons in the ACC we thus recorded from the ACC of 17 rats using silicon probes while they observe another rat receive footshocks (ShockObs condition) and while the implanted animal himself experiences a painful CO_2_ laser (Laser condition). To probe whether neurons are pain selective we also recorded activity while the rats hear a pure tone previously paired with footshocks in a standard fear conditioning paradigm (CS condition) [21] (Fig. 1).

**Figure 1:**
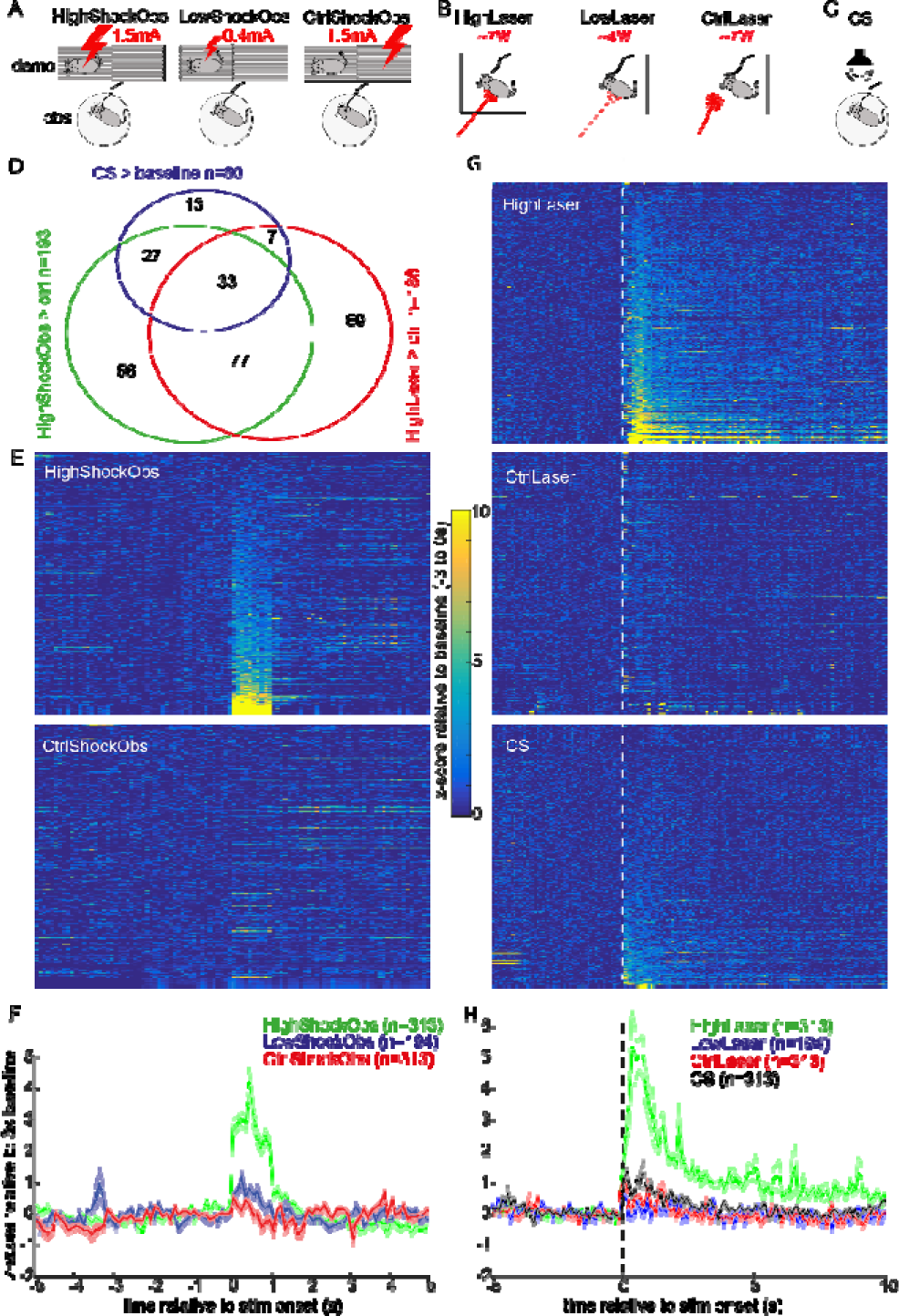
(A) In the ShockObs conditions, the silicon probe implanted animal sits on a circular platform (bottom) while witnessing the demonstrator (top) receive high or low intensity shocks. In the control condition, the shock is delivered to a grid next to the demonstrator and does not trigger pain. The ‘LowShockObs’ conditions were added in the last 10/17 animals only. (B) In the Laser conditions, the implanted animal is alone, and a CO_2_ laser is shone onto his paws or tail. Laser intensity is calibrated individually to trigger pain (HighLaser) r to be just below pain threshold (LowLaser). As a control condition, the laser is shone close to but without touching the animal. (C) In the CS condition, the implanted animal is alon, and a fear conditioned pure tone is played back. (D) Venn diagram specifying the number of MUA channels that show significant responses. Each cell was tested at p<0.01 using a t-test comparing MUA in HighShockObs vs CtrlShockObs (green), HighLaser vs CtrlLaser (red) and CS vs. baseline (blue). Numbers indicate the number of channels that show significant activations in the respective contrast or intersection of contrasts. (E) MUA of the 313 MUA channels tested in High- and CtrlShockObs. Each line shows the z-transformed average MUA response of a channel. Z-transformation was made relative to the mean and std of the 3 seconds prior to each stimulus onset. Stimulus onset is shown as the dotted white line, and the time axis for E and F is shown in F. The channels are ordered as increased average z-score for HighShockObs in the 1s following stimulus onset. (F) Average of E, plus the LowShockObs condition from the n=194 channels acquired in the last 10/17 animals. The shading always represents the standard error of the mean. (G,H) Sa e as E and F for the Laser and CS conditions. For the Laser conditions, channels are ordered in increasing HighLaser responses, in the CS condition in increasing CS response. The x-axis for laser and CS is shown over a longer period because of time due to illustrate the longer MUA response.

## Results

We compared the multiunit activity (MUA[22]) during a baseline period (−1.2s to −0.2s relative to stimulus onset) against the stimulus triggered response (0s to 1s post stimulus onset for Shock and CS, and 0.3s to 1.3s for the Laser response because the latter depends on slower conducting fibers [23]). Stimulus triggered reductions in MUA activity were too rare for further analysis (2/425 following HighShockObs, 6/425 following HighLaser and 3/425 following CS, each tested against their baseline using matched-pair one-tailed t-test at p<0.01). Of the 425 channels that were implanted in the 17 animals, however, 313 (74%) showed increase MUA in at least one condition (matched-pair, one-tailed t-test, HighShockObs>Baseline, HighLaser>Baseline or CS>Baseline, p<0.01, Fig. 1D) and were analysed further. These 313 channels reveal responses to the HighShockObs had short latencies and lasted for about 1s (Fig. 1E-F). Response to the HighLaser, as described in the literature[23], was strong, with a slower onset and lasted for several second (Fig. 1G-H). Responses to the CS were much weaker (Fig 1E-F) despite triggering very robust defensive responses (freezing 69% during the 12 min when CS were played back vs 3.6% in the 12min baseline, t(14)=10.28, p<0.001).

The Venn diagram in Figure 1D shows that there is interesting overlap between channels responding in the different conditions. Of the 313 responsive channels, 62% (193/313) responded significantly more to HighShockObs compared to CtrlShockObs to an empty grid (Fig. 1E-F). Much like in the human ACC, many (71%) of the 193 channels that responded in that social condition also responded when first-hand affective experiences were triggered in the rat using the HighLaser (HighLaser>CtrlLaser) or CS playback (CS>baseline) and will be labelled “mirror channels” hereafter. Most importantly, at the scale of MUA, most of the mirror channels showed selectivity in their response to the animal’s first-hand experience: of the 110 mirror channels responding to HighLaser>CtrlLaser, the majority (77) did not respond to the CS (Fig. 1G-H), and of the 60 mirror channels that responded to CS>Baseline, 27 did not respond to HighLaser>CtrlLaser. Only 33 of the mirror channels responded unselectively to both first-hand conditions. Supplementary Figure 1 illustrates that channels preferring the CS>Laser and those preferring the Laser>CS co-exist in simultaneously recorded channels from individual animals. This illustrates that the selectivity we observe for Laser vs CS in our mirror channels cannot be explained by the animal paying more attention to one condition over another given that on the same trials, some channels preferred one condition and others preferred the other.

In the last 10 of the 17 animals, we added an intermediate intensity of ShockObs and Laser to our experimental design. That the LowLaser condition did not trigger a robust MUA (Fig. 1H) is informative: The LowLaser intensity was chosen to be perceptible but not painful (as judged by the absence of paw licking or similar nocifensive behavior). That ACC channels only respond to the Laser condition once it becomes painful (HighLaser) is compatible with the notion that this ACC response signals pain. The LowShockObs condition, on the other hand, did trigger significant (albeit weaker than HighShockObs) nocifensive behavior in the demonstrator (Fig 2B), and noticeable response in the ACC (Fig. 1F).

**Figure 2.**
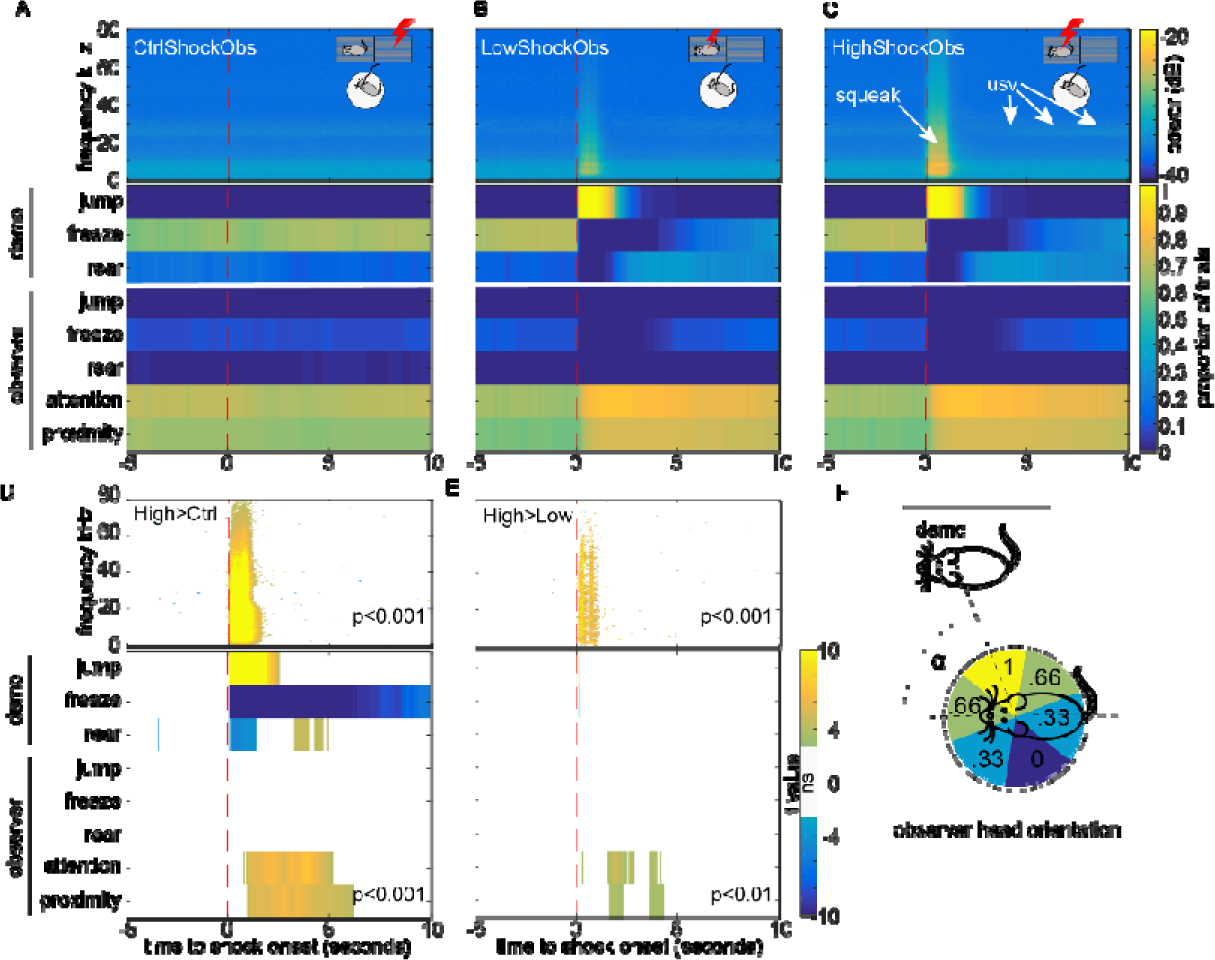
Behavioral scoring of the shock conditions. (A-C) Grand averaged shock-triggered audio spectrogram (top) and ethogram (bottom) obtained by averaging all trials and all animals. Note the broadband signal occurring in the 1s post-stimulus. This includes the pain squeak and the rattling of the cage triggered by the jump. (D) Random effect comparison High>CtrlShockObs done by averaging all the trials per animal, and then using a matched pair-t-test (n=17 animals) pixel per pixel. (E) Same for High>LowShockObs for the 10 animals for which LowShockObs was tested. Tests are thresholded at p<0.001, except for the n=10 animal ethogram comparison in which no difference survives at p<0.001. (F) attention quantified based on the angle α between observer head orientation and demonstrator with ±30º considered maximal (=1), and 180±30º minimal (=0). In the illustrated example, α=70º, and the attention would be scored as 0.66.

**Figure 3.**
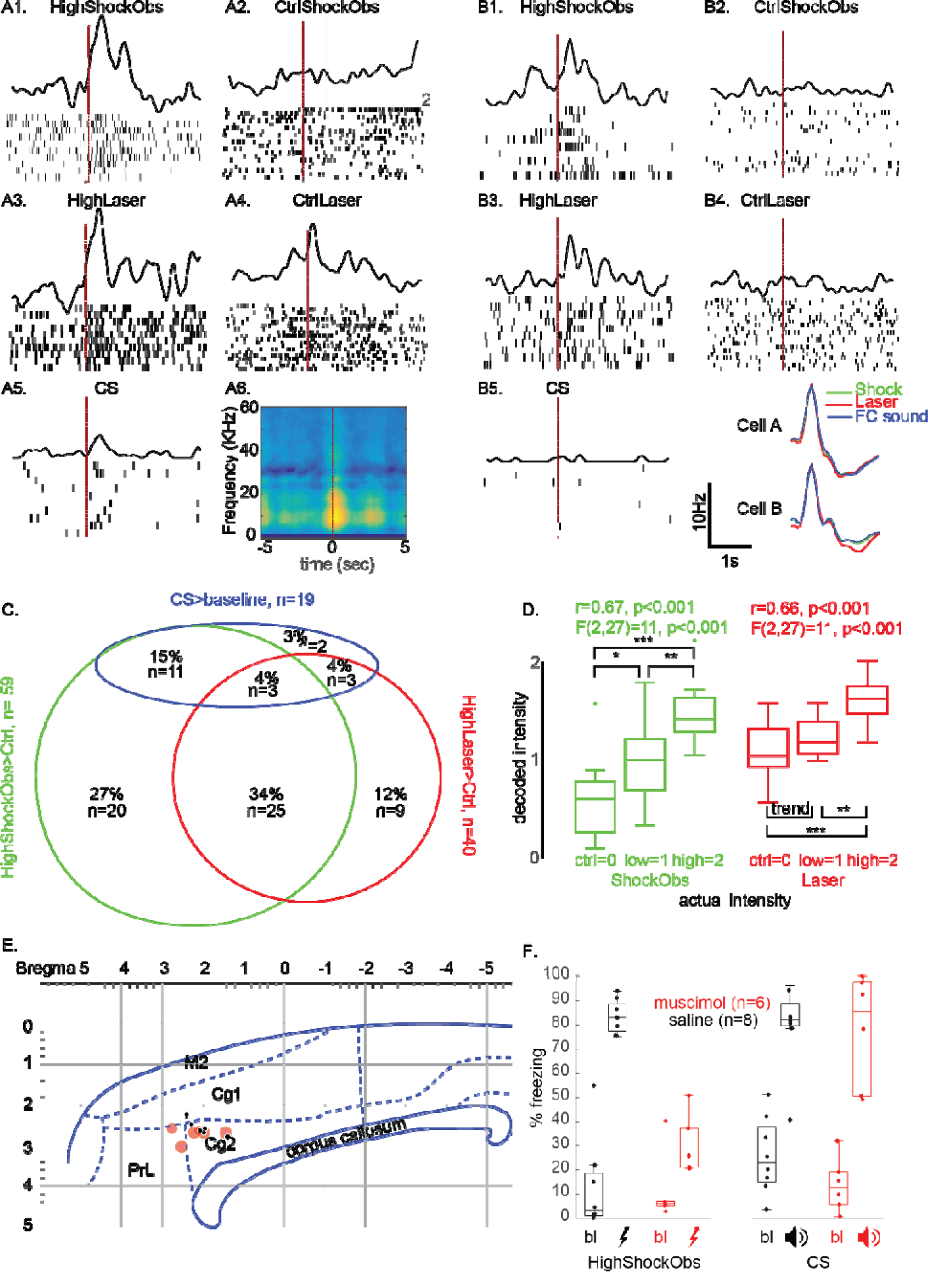
Single unit responses. (A-B) Examples from 2 different animals of cells responding to HighShock>C Obs trl and HighLaser>Ctrl but not CS>Baseline together with average spike shape in each sessions (bottom right). For Cell A, we also show the spike triggered spectrogram in the ShockObs session (A6) evi encing the broadband signature of pain squeaks. The scale bar next to B5 applies to ll spike density functions. (C) Venn dia ram detailing the number of cells showing significant (p<0.05) responses in HighShockObs>Ctrl, HighLaser>Ctrl and CS>Baseline amongst the 73 r sponsive cells. (D) Wisker plot (median & quartiles) of decoded stimulus intensity b sed on an algorithm trained on the Shock bs session and used to either decode leav -one-out ShockObs (green) or Laser (red) spike counts. (E) Locations of the n= muscimol injections (red) and n=8 saline controls (black) on a lateral section. (F) Wisker plot (median & quartiles) of the free ing levels during baseline (bl) or experimental periods (ShockObs or CS playback) in the two groups of animals. *:p<0.05, **:p<0.01, ***:p<0.001. Note that the data rom the ShockObs but not the CS condition is also used to explore how this affects the behavior of the demonstrator in a separate manuscript: https://doi.org/10.1101/452169). In the wisker plot in (D), dots represent outliers, while in (E) the results of each nimal are shown as a dot. All wisker-plots were done using Boxplot in matlab.

Examining the behavior of the demonstrator during the 0-1s interval in which the ACC responded to the HighShockObs condition (Fig. 1F) reveals that this corresponds to the interval in which the demonstrators jumped and squeaked (Fig. 2). This is visible in the spectrogram of the sound-recording as a broadband signal, and in the behavior as a dramatical increase in jumping. The frequency of jumping and intensity of squeaking scaled with Shock amplitude and temporarily interrupts the other behaviors (freezing and rearing). While ultrasonic vocalizations around 22kHz were also apparent following the administration of shocks, their timing was not concentrated on the 0-1s window of HighShockObs ACC response and thus cannot explain the ACC response. The observers’ actions in response to witnessing the shock included turning and walking towards the demonstrator, but this response is delayed by some hundreds of milliseconds after the shock (Fig. 2D) and is thus more likely to represent a downstream effect of ACC activity. The jumping and/or squeaking of the demonstrators seem the most likely trigger of the ACC response to HighShockObs. Importantly, the HighLaser triggered the well described nocifensive reactions to a laser, including paw retraction and licking [24] but did not trigger any squeaking or jumping similar to that in the HighShockObs (Fig. S2). The dual response of some mirror channels to HighShockObs and HighLaser thus cannot be explained by hearing squeaking in both conditions, and must reflect a less trivial association of two physically different stimuli: one signaling the pain of another via exteroception and one signaling potential damage to the animal’s own body via nociceptive afferents.

Spike sorting identified 84 cells spread over 13 animals that could be isolated well and followed over all three experimental sessions. In the remaining 4 animals, low electrode impedance made single cell isolation unreliable. Using the same analysis epochs as for the MUA, amongst these cells, we found 73 responsive cells that showed increased spike counts in at least one condition (HighShockObs>baseline, HighLaser>baseline or CS>baseline, non-parametric Wilcoxon test, p<0.05). Again, there was significant overlap between the responses in the different conditions (Fig. 3A-C). Particularly, 59 (81%) responded to HighShockObs>CtrlShockObs, and of these, 28/59 (47%) also responded to HighLaser>CtrlLaser and 14/59 (24%) to CS>Baseline, thereby demonstrating mirror properties at the single cell level in 66% of the Shock responsive neurons. Most importantly, only 3 of these mirror cells responded to both HighLaser>CtrlLaser and CS>baseline, while all others responded to only one of the first-hand experiences For the majority of the cells (n=25), this took the form of a significant response to HighLaser>CtrlLaser and not to the CS>baseline. Fig. 3A,B illustrates 2 examples of cells from different animals that demonstrate this property. The selectivity of the ACC mirror cells is further borne out by a direct comparison of spike counts for CS and HighLaser in the n=25+3 cells that responded to HighShockObs>CtrlShockObs and HighLaser>CtrlLaser: for 23 of these 28 Shock and Laser cells, HighLaser triggered significantly more spikes than the CS condition (Wilcoxon, p<0.05). This provides the brain with the selectivity necessary to differentiate between states typically labeled as pain (HighLaser) and fear (CS) from the spike count of these neurons. A smaller proportion of mirror neurons seemed selective for CS, with 11 responding significantly to CS>Baseline but not HighLaser> CtrlLaser. Only 3 indiscriminately responded to both CS and HighLaser.

A binomial distribution (59 trials at p=0.05 each) indicates that finding 7 or more amongst the 59 shock responsive cells to respond to another condition is unexpected (p<0.03), and finding 25 pain selective mirror cells is extremely unlikely (p<10^-14^). We therefore found significant evidence for selective emotional mirror properties in the ACC, i.e. that neurons responding to the observation of pain also respond to the experience of pain (HighLaser), but not to other, non-painful salient stimuli (CS). That so few neurons respond to all three conditions (n=3, below what could be expected by chance) points to the fact that the ACC may contain distinct ‘channels’ of neurons mapping another animal’s response to a shock onto the witness’ representations of pain (n=25) or fear (n=11), respectively. Histological reconstruction of the cells showed that mirror cells with different selectivity are intertwined with cells without mirror property along the length of the ACC (Supplementary Figure 3). If the spatial distribution of cells with these properties were similar in humans and rodent, the lack of specificity at the level of fMRI voxels [8,9] may indeed have been the result of pooling the response of neurons with different selectivity within a voxel. To further explore what may trigger the neural response in the HighShockObs condition, we also computed spike-triggered average spectrograms, which revealed the broadband signal typical of pain squeaks to co-occur with moments of high spiking (Fig 3A6).

Using the 69 neurons for which we have 10 trials of the High, Low and Ctrl conditions for ShockObs and Laser, we find that a decoder trained to decode Shock spike counts (Fig. 3D green) successfully decodes Laser spike counts (Fig. 3D, red) with a correlation between actual and decoded stimulus intensity of r=0.66, t(28)=4.7, p<0.001. This suggests that pain observation and pain experience do share a common code, and that the failure of previous attempts to decode pain states across modality from fMRI signals [10] may be due to the poor spatial resolution of fMRI. The pain specificity of the ACC signal is further supported by an ANOVA across the 3 Laser conditions (red, main effect of condition F(2,27)=11, p<0.001) that showed that the two conditions that did not induce pain (CtrlLaser and LowLaser) were decoded as of similar pain intensity (paired t-test p>0.08), whilst the condition that triggered nocifensive behaviour (HighLaser) was decoded as significantly more intense than either of the other two.

Finally, to test if the ACC is necessary to trigger vicarious nocifensive behaviour in the rat, we bilaterally injected muscimol or saline, respectively, into the ACC of two new small groups of witnesses (Fig 3E) and quantified their socially triggered freezing in response to a HighShockObs condition and to a CS playback in separate sessions. In line with previous observations in mice[14], we found that although both groups showed increases in freezing in both HighShockObs and CS sessions relative to their baselines (all t>5.6, all p<0.002), the socially triggered freezing (Shock) was reduced in the muscimol compared to the saline group (t(12)=10.7, p<0.001). This was not true in the non-social condition (CS, t(12)=0.17,p>0.8). The necessity of the ACC for socially rather than non-socially triggered freezing was confirmed by a mixed ANOVA with 2 groups (Saline vs Muscimol) x 2 sessions (Shock vs CS) x 2 Epoch (baseline vs. Shock/CS) that yielded a significant 3 way interaction (F(1,12)=17, p<0.001).

## Discussion

In summary, our data shows that the rat ACC contains mirror-like neurons showing activity increases during Shock observation and Laser experience, and that the population spiking can be used to decode pain intensity across first-hand and witnessed pain. Importantly, for the majority of these neurons, there was evidence for selectivity for the experience of laser-triggered pain over that of other negative affect such as the fear triggered by a CS. Deactivating this region reduces socially triggered freezing without compromising freezing to non-social danger signals (CS).

Although it is difficult to attribute human labels to rodents [25], CS is the prototypical procedure to trigger *fear*, while CO_2_ lasers are a gold standard method for inducing *pain* [26,27]. That many neurons responding to shock observation respond to the laser but not the CS suggests that shock observation may be predominantly mapped onto a representation of *pain* in the self. This dovetails with the fact that the behavioral signature most associated with the response, the squeak, is considered a highly specific pain signal [28]. The vicarious activation of ACC nociceptive neurons may then prime nocifensive behaviors in the observer preparing it to cope with the same source of harm, including orienting towards the danger (Fig. 2) and elevated freezing (Fig 3E) often reported in such paradigms [12,13]. That an, albeit probably smaller, proportion of Shock responsive neurons preferentially respond to the CS suggests that the observer’s ACC may actually map the Shock observation onto a hybrid neural ensemble composed of a majority of pain and a minority of fear representations.

This finding shows that the principle of action-selective mirroring discovered in the motor system while monkeys view or listen to the emotionally neutral actions of others [29,30] also applies to how mammals process the affective signals of others. It is notable, that the brain region in which we find this mechanism [region 24 in ref 19] is similar in location, cytoarchitecture, and connectivity to the location of the human cingulate in which fMRI studies have revealed an increase in BOLD signal during both pain observation and experience [2,19]. If one embraces the notion that mammals may share a common neural mechanism for emotional contagion [1,31] this finding is relevant to the neural basis of human empathy [5,9,32].

## Acknowledgements

We thank Marta Moita for advice on experimental design. We thank Chris de Zeeuw, Cyriel Pennarz and Francesco Battaglia for methodological advice and help. We thank J. Hernandez-Lallement, R. Rajendran, E. Soyman and S. Voges for critical comments on the manuscript.

## Author Contribution

CK and VG conceived the study and acquired the funding. All authors helped refine the final experimental design. MC & YH conducted the experiments with help of FM and ML. MC, YH and CK analyzed the data with critical comments from VG. CK wrote the manuscript with help from VG, and all authors commented on the manuscript.

## Data and Code availability

Data and Code will be made available upon reasonable request.

## Supplementary Figures

**Figure S1:**
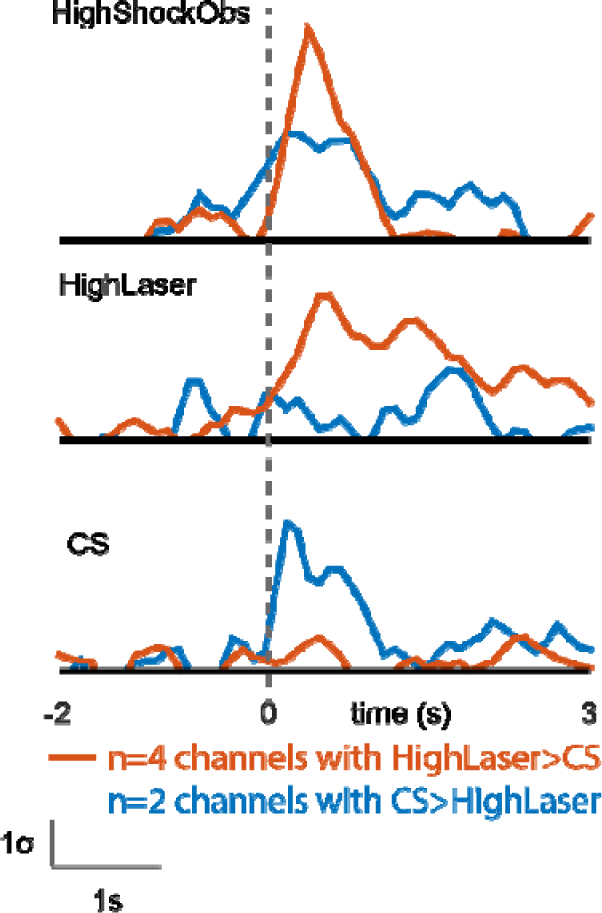
Opposite preference in simultaneously recorded MUA channels. The figure represents the average MUA response of two small populations of MUA channels that were recorded simultaneously in one animal (#31) and had opposite preferences for Laser vs. CS. The red line represents the average of 4 channels that responded to HighLaser>CtrlLaser (p<0.01 for each channel) but not CS>Baseline (p>0.2 for each channel). The blue line represents the average of 2 channels that responded to CS>Baseline (all p<0.01) but not HighLaser>CtrlLaser (all p>0.2). All 6 channels showed significant responses to ShockObs (HighShockObs>CtrlShockObs, all p<0.01). The time courses illustrate how within the same animal, simultaneously recorded channels can share a sensitivity to the signals of another rat (top pannel) but show opposite selectivity during the experience of HighLaser (red) and CS (blue). Given that they were recorded simultaneously, this cross-over cannot be explained by differences in salience of or attention to Laser vs. CS.

**Figure S2:**
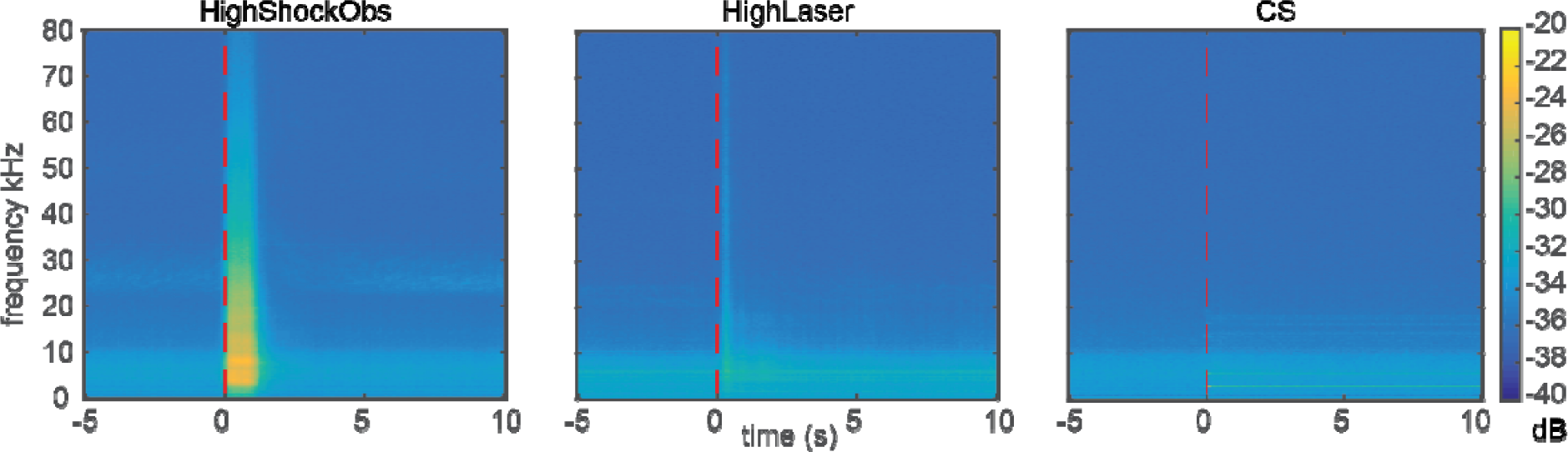
Average Spectrogram. The three conditions were accompanied by quite different soundscapes. While the HighShockObs condition was characterized by squeaking in the audible range (left), the HighLaser was characterized by a subtle sound resulting from the rat’s bodily movements (e.g. paw retraction). The CS finally shows the signature of the 8kHz CS and its harmonics.

**Figure S3:**
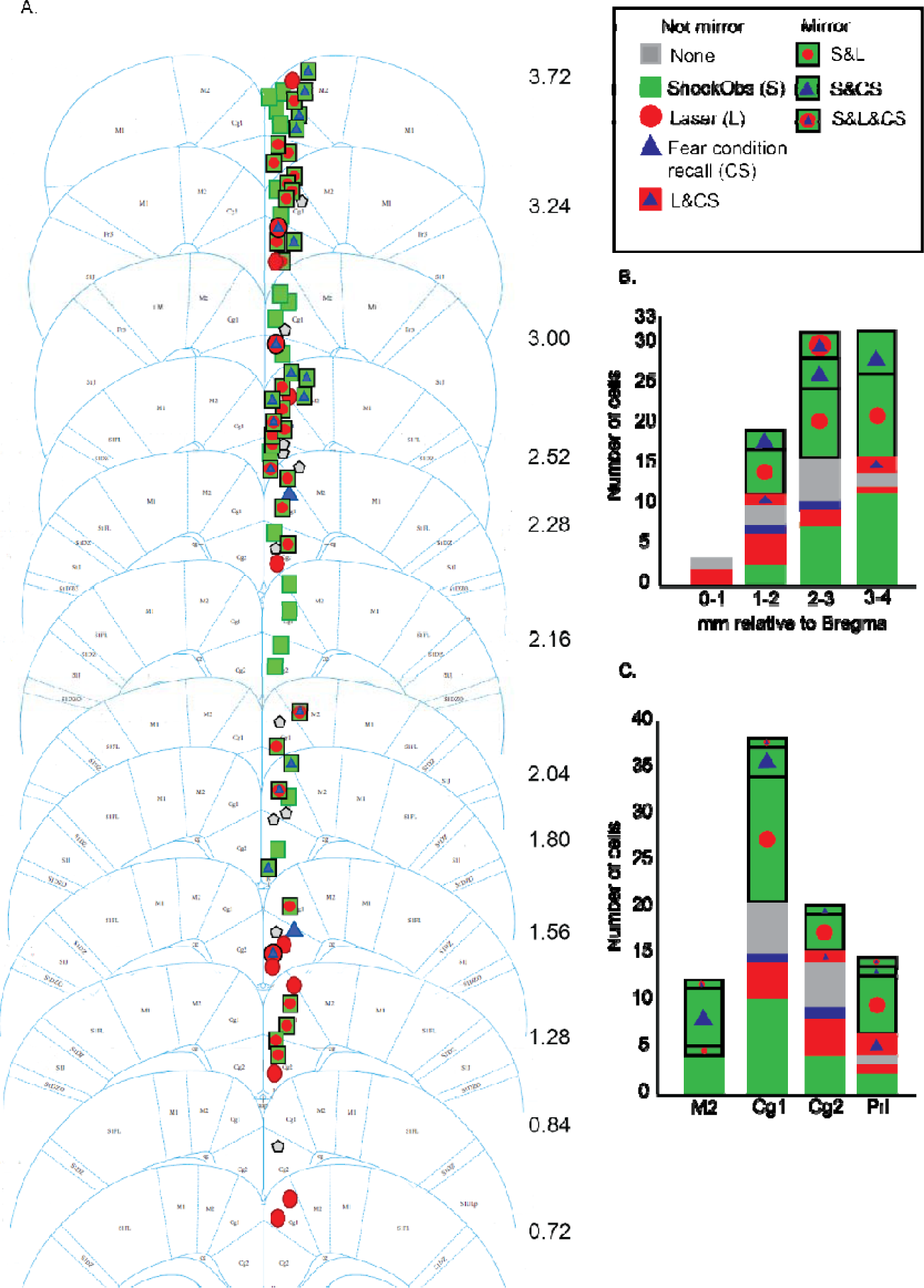
Histological reconstruction of cells with different properties. (**A)** Reconstruction of the isolated cells on coronal slices. The Bregma AP (Anterior-Posterior) coordinate is indicated in mm next to each slice from the Paxinos Atlas[33]. (**B)** Number of cells of each category classified by their AP coordinate. Although we isolated more cells in more anterior parts of the ACC, a chi^2^ test comparing the proportion of mirror cells (i.e. Shock&Laser, Shock&CS and Shock&CS&Laser) across the 4 AP bins reveals that there was no significant difference in proportion as a function of AP coordinates (all p>0.2). (**C)** Cell counts as a function of cytoarchitectonic division. Most of the cells we isolated in Cg1 and Cg2, corresponding to Area 24, however the proportion of mirror neurons was similar across these regions (Chi^2^ all p>0.05).

## Supplementary material and methods

### Subjects for the electrophysiological experiments

34 male long Evan rats (6-8weeks old/250-350g) were obtained from Janvier, France for the electrophysiological experiments (see section *Muscimol Experiment* for all methods regarding that additional experiment). Animals were randomly assigned to different roles, consisting of 17 observers and 17 shock demonstrators. Upon arrival all animals were socially housed at ambient room temperature (22-24 °C, 55% relative humidity, SPF), on a reversed 12:12 light:dark cycle (lights off at 07:00). Food and water was provided *ad libitum*. All experimental procedures were pre-approved by the Centrale Commissie Dierproeven of the Netherlands (AVD801002015105) and by the welfare body of the Netherlands institute for Neuroscience (IVD, protocol number NIN161107).

### Test setups

All habituations and tests were conducted inside a faraday cage, in dim red light during the dark part of the circadian rhythm, with background radio turned on. Three different set-ups were used based on condition (Fig. S4). During the shock observation test (Fig. S4A), the observers were placed in an elevated circular platform (H:70cm, 30 cm diameter), surrounded by a transparent plastic wall of 2 cm in height, and with bedding from the observer s home cage. The demonstrator’s testing box consisted of two chambers separated by a perforated Plexiglas divider (each: L24cm xW:25cm x H:34cm) with stainless still grid floors. The cage was positioned close to the observer platform with the chamber containing the demonstrator closest to the observer to ensure it was clearly visible to the observer. The wall facing the observer’s platform was made of fine wire mesh (Med associates Inc, USA). During the fear conditioning recall test (CS, Fig. S4C), the cage of the demonstrator was removed, and the observer was placed in the same elevated platform as that used for shock observation, with a buzzer-like loudspeaker playing the conditioned cue (CS) placed ∼30cm away from the platform. Lastly, the observer experience of the heat laser (Laser, Fig. S4B) was conducted on a rectangular stainless-steel metallic platform (15cmx15cm), elevated 30cm and with a 0.5cm fence. The CO_2_ laser was placed outside the faraday cage and the arm used for delivering the heat pulses protruded into the faraday cage, with its tip 15cm away from the observer’s platform. During all tests, behavior, vocalizations and neural activity were recorded using a top and side video camera (Basler acA1300 and mediarecorder software, Noldus, Netherlands), a condenser ultrasound microphone (Avisoft-bioacustics, CM16/CMPA, Germany) and an electrophysiology acquisition system (digital lynx SX and cheetah software, Neuralynx, USA), respectively. To avoid contextual fear, the test pre-exposure was done in a two compartment cage that was different from that used in any of the electrophysiological testing: a two chamber box with angled soft plastic walls (each: L31cm xW:24cm (bottom) L40xW:31(top) x H:44cm) separated by a perforated Plexiglas divider with stainless still grid floors. This pre-exposure box was also washed with a differently scented soap, the background radio was turned off, and light intensity was higher to prevent generalization across sessions.

**Figure S4:**
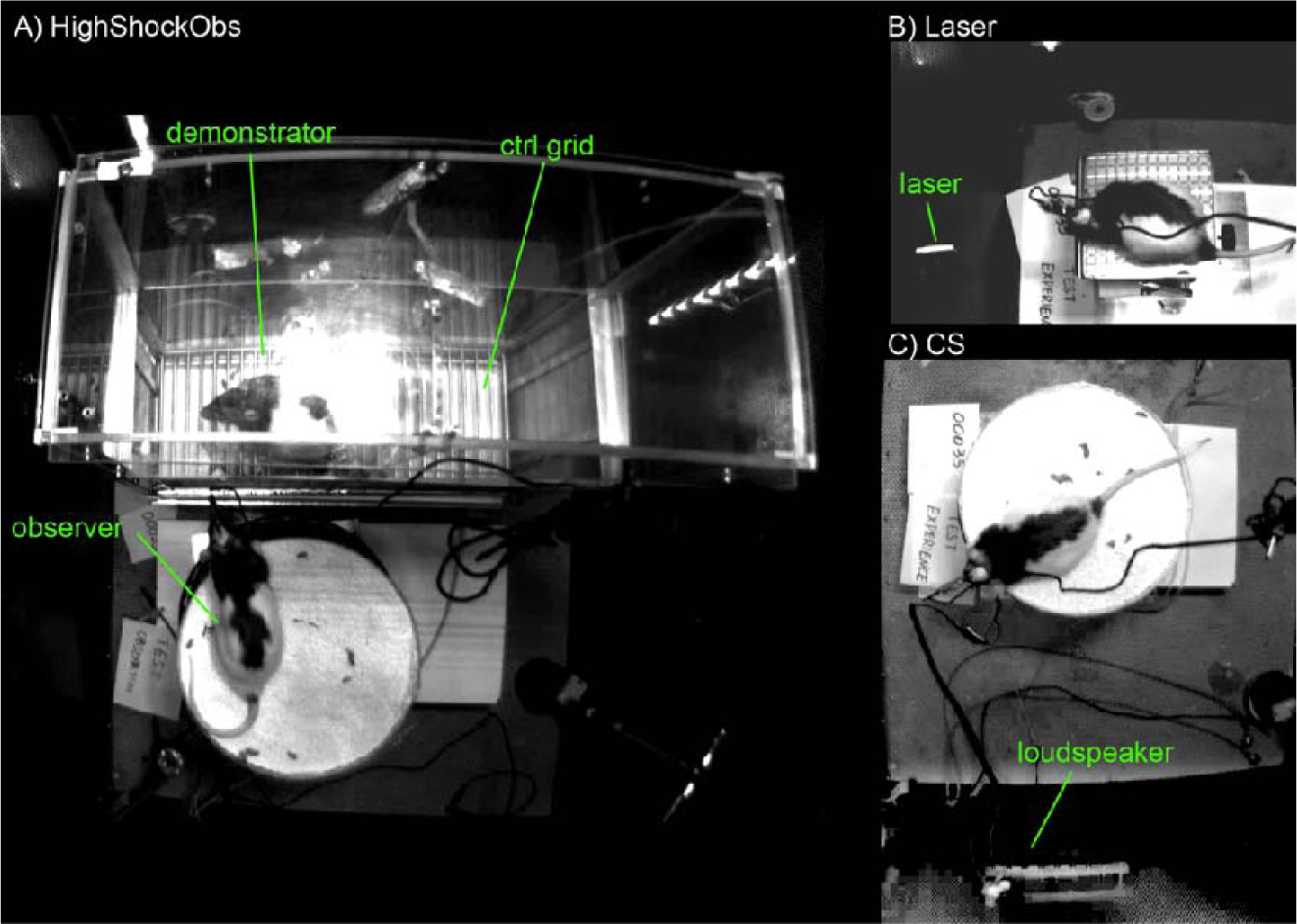
snapshots from the overhead camera illustrating the setups for the three experimental conditions.

### Experimental procedures

#### Acclimatization and pre-exposure

Upon arrival, animals were allowed to acclimate to the colony room for 7 days (week 1, Fig. S5A). To reduce stress and habituate animals to the researchers, during the second week (week 2) all animals were handled every other day for 3 minutes per day. During week 3, to prepare and familiarize the observer with the conditions they will encounter during the test days (ShockObs, CS, and Laser), the observers experienced three types of stimuli: footshocks, fear conditioninging and CO_2_ heat laser. The footshocks and fear conditioning were combined into a single pre-exposure session. The observer animal was put into one compartment of the above-descrined pre-exposure box and a 10 minute baseline was followed by the presentation of five 20s tones (8KHz, 70dB), each associated with the delivery of a 1s shock (0.8mA) during the last second of the tone presentation (1 s at 0.8mA with 60 secs inter-shock interval; shocker model ENV-414 from Med associates, Inc). To prepare the observer for the laser condition it was important to first measure the pain threshold for each animal, which was determined as the stimulation level at which the animal showed consistent paw retraction and/or licking. A laser pre-exposure session was then performed, in which, after a 2 minute baseline, a CO_2_ heat laser (CL15 model:M3) was used to deliver 5 pulses (wavelength 10.6µm, 200msecs, at 60-70% of the total laser power of 15W, beam diameter <2mm) aimed at the paws or tail, with a random inter-stimulus interval of 24 to 36 secs. Shock demonstrators were left in their home cage ensuring they will be naïve to the stimuli on test day.

**Figure S5:**
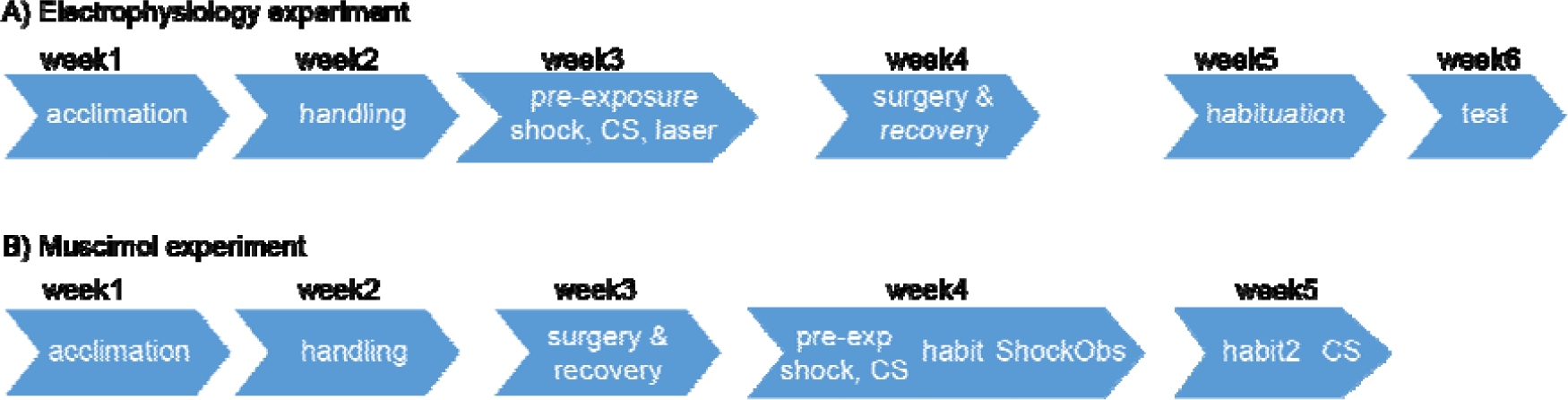
Timing of experimental procedures.

#### Surgery

On week 4, observers underwent a surgical procedure for the unilateral implantation of a multi-shank silicon probes (Atlas Neuroengineering, Belgium, E32-400-SSL4-500), targeting the right anterior cingulate cortex (ACC). Buprenorphine was used for pain relief (30 mins prior to surgery, s.c. 0.01-0.05mg/kg) and isoflurane as anesthetic (4% for induction and 0.8-2% for maintenance). Body temperature and other physiological parameters were monitored throughout the surgery. Once animals were deeply anesthetized, they were placed in a stereotaxic apparatus, six screws were attached to the skull (two of them used to connect the ground wires), a craniotomy was performed (4-4.5mm in diameter) and then the probe was lowered to the target area (centered at Bregma AP:0.96mm, ML:0.3mm, DV:-3mm, angle from vertical: 20o) and secured using multipurpose cement (GC Fuji PLUS capsules, GC Europe N.V., Belgium). After termination of the surgery, animals received Atipamezole (0.5mg, ip) and Metacam (1mg/kg) and were placed in an incubator until they woke up. Thereafter, they were placed in a modified homecage and received wet food until behaviour and weight was recovered. Behaviour, state of the incision and weight was monitored daily for 3 days and once a week thereafter. Animals were allowed to recuperate for at least 7 days prior to test start.

For the rest of week 4 and 5, observers and demonstrators were habituated to the experimental setups for five days (20 minutes/day/setup). On the last three habituation sessions, observers were tethered to the electrophysiology recording system.

#### Electrophysiology

On week 6, the electrophysiological recording sessions were conducted during two consecutive days. Day one included Shock observation followed by Laser experience and on day two, the CS recall test. Separation of the tests onto two different days was done to ensure that baseline activity in the ACC during the second aversive experience does not reflect a carry-over from the previous first-hand experience. Laser preceded the CS session because the question of whether the same cells respond to pain observation and experience was primary, and the question of selectivity with regard to CS secondary. That fewer cells respond to CS than Laser should therefore be interpreted cautiously.

*Shock observation:* test started with a 12-minute baseline, followed by the observer witnessing the demonstrator experience different shock intensities. In the first 7 animals we tested two conditions: 10 high intensity (1.5mA) 1s shocks and 10 control 1s shocks delivered to a grid in the compartment adjacent to that of the demonstrator (intershock interval 60 or 90 secs). In the last 10 animals we added 10 additional 1s shocks of low intensity (0.4mA) and increased the number of control shocks to 20. 0.4mA was chosen because this still leads to a visible and audible reaction of the demonstrator but one that is clearly less intense than that at 1.5mA (Fig. 2).

*Laser experience* test was conducted at least 20 mins after shock observation and it consisted of a 5 minute baseline followed by the laser stimulation trials. As for Shock, in the first 7 animals, we tested 2 conditions consisting of 10 high intensity stimulations (50-70% of the laser power depending on the pre-determined pain threshold per animal), and 10 control stimulations targeting the air near the animal. Each laser stimulation lasted 200ms, and were separated by an inter-laser interval of either 24 or 36 s. In the last 10 animals we added an intermediate intensity for a total of 40 trials: 10 high intensity stimulations, 10 low intensity stimulations (20% less than the HighLaser intensity), and 20 control stimulations. In the high and low intensity trials, the paw or tail of the animal were targeted. Using 20% less laser intensity was chosen to have a more restrictive control stimulus in which the laser is shone onto the animal inducing a termal sensation without inducing pain. The lack of clear nocifensive behavior (no paw licking or retraction) supports the absence of pain at the chosen low intensity.

*CS recall* session consisted of a 12 minutes baseline and 12 minute test period, in which the CS tone (8kHz, 70dB; pre-conditioned with footshocks) was presented 10 times for 20 seconds each time. The inter-CS interval was either 60 or 90 seconds in a pseudorandomized order.

#### Histology

After completion of the experiment, animals were deeply anesthetized and an electrical lesion was performed to mark the positions of the electrodes (2µA, 10 secs, across the top and bottom most contact of each leg). Animals were then intracardially perfused with phosphate buffered saline (PBS, 7.4pH) followed by 4% paraformaldehyde, brains were removed, cut with a cryostat (50µm coronal sections), and Nissl stained for verification of shafts track and electrode positioning.

### Data acquisition

The electrophysiology signals (continuous and single units) were unit gain amplified using a head stage pre-amplifier (HS-36, Neuralynx), relayed to an input board differential input amplifiers with a gain of 15 and acquired with a sampling frequency of 32kHz. For single units, the signal was bandpass filtered (0.6-6 kHz), timestamped and recorded for 1 msec every time the signal passed a manually set threshold.

### Data analysis

#### Behavioral data

The onset and duration of the observer and demonstrator behavior during the tests were manually scored offline in a continuous manner using the open source Behavioral Observation Research Interactive Software (BORIS, Friard & Gamba, 2016)and analyzed using Matlab (MathWorks Inc., USA). For the shock observation test, freezing, rearing, head location, jumping and head orientation of the observer and demonstrator were scored. The observer’s head orientation relative to the line connecting the heads of the two animals was used to quantify attention (Figure 2F), while the distance between the head of the observer and the demonstrator’s cabinet was used to measure proximity. Soundtracks of the session were also analyzed using Matlab to extract power per frequency (0-80kHz) relative to the onsets of each condition (i.e. HighShockObs, low shock & control trials; Figure S2). Specifically, the time-frequency decomposition was averaged over all trials of a given condition, and conditions were then compared by performing a t-test separately at each time and frequency by including one value per animal akin to the random effect mass-univariate analyses typical of neuroimaging data. For the laser experience test, the reaction of the observer to the stimuli was scored offline to confirm successful laser targeting. Trials with no behavioral reaction to HighLasers were excluded from further analyses. For the CS recall test, sleeping, grooming and freezing were scored offline to verify classic fear response to the CS.

#### Multiunit activity

Multiunit activity analyses were performed using the FieldTrip toolbox (Oostenveld et al., 2011) and custom-made Matlab scripts. Data from the 32 contacts were first visually explored to identify artifacts. This was done (a) taking the raw signal from each of the 30 high impedance contacts relative to the low impedance reference channel, high-pass filtering it at 1000 Hz, rectifying and low-pass filtering at 200 Hz to approximate MUA. Trials in which extreme MUA activity (z>8 or z<-5) occurred across many or all channels were removed. This lead to the rejection of 7 trials in total. Inspection of the video recordings identified 5 more trials that had to be rejected because the Faraday cage had to be opened for experimental reasons. We then performed a pair-wise re-referencing of the cleaned data on each electrode shaft offline. Each of the 5 shafts of the electrode had 6 contacts, and re-referencing was done by subtracting raw data from the vertically adjacent contact of each leg resulting in 5 channels (each the difference of two contacts) per shaft, or 25 channels per animal. The data was then high-pass filtered at 1000 Hz, rectified and low-pass filtered at 200 Hz as recommended in [22]. Trials were created 2 seconds before to 3 seconds after the onset of the stimuli.

For statistical analyses, we used the surface under the MUA. Specifically, we computed the area under the MUA during the baseline epoch (−1.2s to 0.2s relative to stimulus onset) and during an epoch of same duration (1s) during stimulus presentation when we expect the response to occur. For Shock and CS trials, we expect low latency responses and therefore used the epoch 0s to 1s post stimulus onset. For Laser, three reasons lead us to expect longer latencies: (a) the laser pulse had a 200ms duration and the thermal energy does not reach its maximum (and hence painful level) before 200ms, (b) the burning pain most associated with affective reactions depend on unmyelinated fibers with slow conduction times and (c) laser evoked spiking in the ACC has been reported to reflect laser intensity reliably from ∼300ms after stimulus onset [23,34]. Accordingly, for that condition we shifted the experimental epoch to 0.3s to 1.3s post laser onset. The same duration interval was used in all conditions (1s) to avoid biasing analyses for a particular condition. One tailed t-tests were then used to compare responses against baseline or across conditions because the surface under the MUA in 1s intervals were continuously and approximately normally distributed. Paired t-test were used when comparing an experimental period against its directly preceding baseline, and two sample t-test were used when comparing the experimental period across conditions. The significance threshold was set at 0.01.

We first identified channels that were responsive by requiring that HighLaser > baseline OR HighShockObs > baseline OR CS > Baseline. We then explored the selective responsivity of those channels that were ‘responsive’ by comparing conditions against their control condition (i.e. HighLaser > CtrlLaser; HighShockObs > CtrlShockObs). For the CS playback we were more lenient in order to detect all cells that respond to any salient sound, and thus compared CS against baseline (CS>baseline).

#### Single unit data

To characterize the response of units that respond to witnessing a shock to another animal, the data from the shock observation session were clustered using spikesort 3D (Neuralynx) and KlustaKwik (Harris et al., 2000) then manually examined to identify the channels in which there were single units present. Channels without single unit activity were excluded from further processing. To ensure that the same single unit was present across all sessions (i.e., Shock observation, Laser experience and CS recall) the data from each contact from the three sessions was merged into a single file and then processed and analyzed as if it was a unique session. This merged data set was then automatically clustered and manually cleaned using SpikeSort 3D and KlustaKwik. To further ensure that the same cell was present across all sessions, spike waveforms and firing rate during baseline period of the Laser and CS recall sessions were compared to the shock session and a cleanup procedure was conducted as follows: 1) spikes with a waveform-correlation lower than 0.85 with the average shock waveform were removed, 2) spikes with a peak amplitude beyond ±15% of the peak amplitude of the average shock waveform were removed and 3) sessions with a firing rate ratio between the Shock baseline period and the other sessions’ baseline period higher than 8 were removed (after steps 1 and 2 had been applied). In addition, sessions with a spike firing rate lower than 0.06Hz were not included in the final analysis.

Statistical analyses were performed on spike counts from epochs defined as for the MUA analysis: baseline epoch were always from −1.2 to −0.2s relative to stimulus onset, and experimental epochs were from 0 to 1s for Shock and CS, and 0.3 to 1.3s for Laser. Spike counts were compared against their baseline using one-tailed Wilcoxon signed-rank test, threshold set at *p*<0.05. This non-parametric test was used instead of a t-test at p<0.01 because spike counts are discrete numbers that are sometimes ill distributed. However analyzing the same data with t-tests at p<0.01 lead to conceptually similar results. Spike counts were compared across conditions using the Wilcoxon rank sum test at p<0.05. Clusters were classified as responsive in a given session, if HighShockObs > Shock Baseline OR HighLaser > Laser Baseline OR CS > CS Baseline. For shock and laser sessions, clusters were classified as specific in their response in a given condition if HighShockObs > CtrlShockObs or HighLaser > CtrlLaser. As for the MUA, CS responses were simply assessed compared to baseline to be sensitive in detecting any response to a salient event.

*Spike triggered spectrogram:* To further characterize the features triggering the activity of interesting neurons (Fig. 3A6) in the Shock session we performed spike triggered analyses of the sound recording. For this analysis, we did not consider the first 12 minutes of baseline but focused on the continous period starting shortly before the first shock and ending after the last shock. In that period, instantaneous firing rate of each neuron was calculated as the inverse of the inter-spike interval. Moments of unusual firing were then identified as moments with instantaneous firing rate in the top 5% of all instantaneous firing rates. For each of these moments, we then extracted the 5s prior and after the surprising spike rate and averaged this spectrogram across all surprising spike rates. Finally, to identify *changes* of power associated with the spikes, we subtract for each moment at a given frequency the average power in that frequency across the whole experimental period.

#### Muscimol experiment

*Note*: data from the ShockObs condition of the muscimol experiment described here is also used to explore the effect this has on the behavior of the demonstrator in another manuscript currently under revision at Plos Biology (see https://doi.org/10.1101/452169 for a preprint).

*Subjects*: 48 male long Evan rats (6-8weeks old/250-350g) were obtained from Janvier, France. Animals were randomly assigned to two groups: saline control group (n=24, 12 observers and 12 demonstrators) or muscimol group (n=24,12 observers and 12 demonstrators). Of those, 8 saline and 6 muscimol observers could be included in the final analysis after removing those with damaged or clogged canulae and those in which histological reconstruction revealed damage to the corpus callosum (see below). Upon arrival all animals were housed in observer-demonstrator dyads at ambient room temperature (22-24 °C, 55% relative humidity, SPF), on a reversed 12:12 light:dark cycle (lights off at 07:00). Food and water was provided *ad libitum*. All experimental procedures were pre-approved by the Centrale Commissie Dierproeven of the Netherlands (AVD801002015105) and by the welfare body of the Netherlands institute for Neuroscience (IVD, protocol number NIN151104).

#### Habituation and Surgery

on Week1 (Fig. S5B) after arrival, animals were acclimated to the colony room for 7 days. On Week2 animals were handled every other day for 3 minutes. Thereafter, cannulae were implanted into the ACC. All animals were anesthetized by isoflurane. The animals were then positioned in a stereotaxic frame with blunt-tipped ear bars, and a midline incision was made. Six holes were drilled (2 for anchoring screws and 1 for the cannula per hemisphere). Two single guide-cannulas (62001; RWD Life Science Co., Ltd) were implanted targeting bilateral ACC (AP, +1.7; ML, ±1.6; DV, +3.5 mm with a 20° angle from the surface of the skull, Paxinos and Watson, 1998) and chronically attached in the observer animals with a thin layer of acrylic cement (Super-Bond C & B ®, Sun Medical Co. Ltd., Shiga, Japan) and thick layers of acrylic cement (Simplex Rapid, Kemdent, UK). To prevent clogging of the guide cannula, a dummy cannula (62101; RWD Life Science Co., Ltd) was inserted and secured until the microinjection was administered. After a week of recovery, observers were habituated to fake micro-infusion and to the experimental setup for the HighShockObs condition with their demonstrator for 20 minutes. The testing box consisted of two chambers (each: L24cm xW:25cm x H:34cm) divided by a perforated Plexiglas divider, with a stainless-steel grid as a floor on the side of the demonstrator and a plastic platform on the side of the observer (Med associates Inc, USA). The observer then was pre-exposed to footshocks using the same protocol described in the electrophysiology experiment above. Demonstrators were naïve to the shock and tone stimuli.

#### HighShockObs

three days after pre-exposure the HighShockObs test was performed. Fifteen minutes prior to the shock observation test, observer animals were lightly restrained, the stylet was removed and an injection cannula (62201; RWD Life Science Co., Ltd) extending 0.8 mm below the guide cannula was inserted. Muscimol (0.1 μg/μl) or saline (0.9%) was microinjected using a 10 μl syringe (Hamilton), which was attached to the injection cannula by PE 20 tubing (BTPE-20; Instech Laboratories, Inc.). A volume of 0.5 μl per side was injected using a syringe pump (70-3007D; Harvard Apparatus Co.) over a 60 s period, and the injection cannula remained untouched for an additional 60 s to allow for absorption into the brain region and to minimize injectate along the track of the cannula. The protective cap was secured to the observer animal after the infusion and then the animal was returned to the home cage. Six (2 from saline and 4 from muscimol group) were excluded due to damaged or clogged cannulas. Shock observation test then started with a 12-minute baseline, followed by the observer witnessing the demonstrator experience 5 footshocks (1sec, 1.5mA each, pseudorandom intershock interval 120 or 1800 secs).

#### CS

One week later, the observers underwent the conditioned stimulus recall test. Fifteen minutes prior to the test, microinjection of muscimol or saline were performed with the same protocol as prior to the shock observation test. Observers were then put into a skinner box in a context that was different from the shock observation test (i.e. different smell, illumination, and floor texture). After 12 minutes baseline, the CS tone were played for 5 times (20 secs each, 120 or 180 secs pseudorandom interval). All test sessions were videotaped using a Basler GigE camera (acA1300-60gm) controlled by MediaRecorder 2 (Noldus, the Netherlands).

#### Histology

After completion of the experiment, animals were intracardially perfused with phosphate buffered saline (PBS, 7.4pH) followed by 4% paraformaldehyde, brains were removed, cut with a cryostat (50µm coronal sections) and Nissl stained for verification of cannula location. Four dyads (2 from saline group and 2 from muscimol group) were excluded from data analyses after histology examination suggesting damage of corpus callosum due to injection.

#### Data analysis

The freezing of the observers and demonstrators were scored as in the electrophysiology experiment. Freezing time was calculated as the sum of all freezing moments in a certain epoch and freezing percentage was calculated as the total freezing time divided by the total time of the epoch. Baseline period was defined as the first 720 seconds of the test and the test period was defined as the 720 seconds after baseline. For comparison between periods (baseline vs shock/CS) and conditions (muscimol vs control), repeated measures ANOVAs (IBMSPSS statistics, USA) were performed with baseline and test period as within subject factors and the conditions as between-subject factors.

## Bibliography

1. de Waal, F.B.M., and Preston, S.D. (2017). Mammalian empathy: behavioural manifestations and neural basis. Nat. Rev. Neurosci. 18, 498–509. Available at: http://www.ncbi.nlm.nih.gov/pubmed/28655877 [Accessed September 7, 2017].

2. Lamm, C., Decety, J., and Singer, T. (2011). Meta-analytic evidence for common and distinct neural networks associated with directly experienced pain and empathy for pain. Neuroimage 54, 2492–2502.

3. Singer, T., Seymour, B., O’Doherty, J., Kaube, H., Dolan, R.J., and Frith, C.D. (2004). Empathy for pain involves the affective but not sensory components of pain. Science 303, 1157–62. Available at: http://www.ncbi.nlm.nih.gov/pubmed/14976305 [Accessed July 10, 2014].

4. Meffert, H., Gazzola, V., den Boer, J.A., Bartels, A.A.J., and Keysers, C. (2013). Reduced spontaneous but relatively normal deliberate vicarious representations in psychopathy. Brain 136, 2550–62. Available at: http://www.pubmedcentral.nih.gov/articlerender.fcgi?artid=3722356&tool=pmcentrez&renderty pe=abstract [Accessed January 7, 2016].

5. Gallese, V., Keysers, C., and Rizzolatti, G. (2004). A unifying view of the basis of social cognition. Trends Cogn. Sci. 8, 396–403. Available at: http://www.ncbi.nlm.nih.gov/pubmed/15350240 [Accessed July 10, 2014].

6. Hutchison, W.D., Davis, K.D., Lozano, A.M., Tasker, R.R., and Dostrovsky, J.O. (1999). Painrelated neurons in the human cingulate cortex. Nat. Neurosci. 2, 403–405. Available at: http://www.ncbi.nlm.nih.gov/pubmed/10321241 [Accessed January 11, 2018].

7. Sakaguchi, T., Iwasaki, S., Okada, M., Okamoto, K., and Ikegaya, Y. (2018). Ethanol facilitates socially evoked memory recall in mice by recruiting pain-sensitive anterior cingulate cortical neurons. Nat. Commun. 9, 3526. Available at: http://www.nature.com/articles/s41467-018- 05894-y [Accessed September 30, 2018].

8. Wager, T.D., Atlas, L.Y., Botvinick, M.M., Chang, L.J., Coghill, R.C., Davis, K.D., Iannetti, G.D., Poldrack, R.A., Shackman, A.J., and Yarkoni, T. (2016). Pain in the ACC? Proc. Natl. Acad. Sci. 113, E2474–E2475. Available at: http://www.ncbi.nlm.nih.gov/pubmed/27095849 [Accessed September 30, 2018].

9. Zaki, J., Wager, T.D., Singer, T., Keysers, C., and Gazzola, V. (2016). The Anatomy of Suffering: Understanding the Relationship between Nociceptive and Empathic Pain. Trends Cogn. Sci. 20, 249–59. Available at: http://linkinghub.elsevier.com/retrieve/pii/S1364661316000449 [Accessed March 9, 2016].

10. Krishnan, A., Woo, C.-W., Chang, L.J., Ruzic, L., Gu, X., López-Solà, M., Jackson, P.L., Pujol, J., Fan, J., and Wager, T.D. (2016). Somatic and vicarious pain are represented by dissociable multivariate brain patterns. Elife 5. Available at: http://www.ncbi.nlm.nih.gov/pubmed/27296895 [Accessed June 14, 2018].

11. Corradi-Dell’Acqua, C., Tusche, A., Vuilleumier, P., and Singer, T. (2016). Cross-modal representations of first-hand and vicarious pain, disgust and fairness in insular and cingulate cortex. Nat. Commun. 7, 10904. Available at: http://www.ncbi.nlm.nih.gov/pubmed/26988654 [Accessed September 17, 2018].

12. Atsak, P., Orre, M., Bakker, P., Cerliani, L., Roozendaal, B., Gazzola, V., Moita, M., and Keysers, C. (2011). Experience modulates vicarious freezing in rats: a model for empathy. PLoS One 6, e21855. Available at: http://www.pubmedcentral.nih.gov/articlerender.fcgi?artid=3135600&tool=pmcentrez&renderty pe=abstract [Accessed December 6, 2015].

13. Carrillo, M., Migliorati, F., Bruls, R., Han, Y., Heinemans, M., Pruis, I., Gazzola, V., and Keysers, C. (2015). Repeated Witnessing of Conspecifics in Pain: Effects on Emotional Contagion. PLoS One 10, e0136979. Available at: http://www.pubmedcentral.nih.gov/articlerender.fcgi?artid=4565705&tool=pmcentrez&renderty pe=abstract [Accessed January 12, 2016].

14. Jeon, D., Kim, S., Chetana, M., Jo, D., Ruley, H.E., Lin, S.-Y., Rabah, D., Kinet, J.-P., and Shin, H.-S. (2010). Observational fear learning involves affective pain system and Cav1.2 Ca2+ channels in ACC. Nat. Neurosci. 13, 482–8. Available at: http://www.nature.com/articles/nn.2504 [Accessed September 12, 2018].

15. Gonzalez-Liencres, C., Juckel, G., Tas, C., Friebe, A., and Brüne, M. (2014). Emotional contagion in mice: The role of familiarity. Behav. Brain Res. 263, 16–21. Available at: http://dx.doi.org/10.1016/j.bbr.2014.01.020.

16. Keum, S., and Shin, H.-S. (2016). Rodent models for studying empathy. Neurobiol. Learn. Mem. 135, 22–26. Available at: http://www.ncbi.nlm.nih.gov/pubmed/27475995%5Cnwhttp://linkinghub.elsevier.com/retrieve/pii/S1074742716301198.

17. Kim, S., Matyas, F., Lee, S., Acsady, L., and Shin, H.-S. (2012). Lateralization of observational fear learning at the cortical but not thalamic level in mice. Proc. Natl. Acad. Sci. 109, 15497–15501. Available at: http://www.ncbi.nlm.nih.gov/pubmed/22949656 [Accessed September 12, 2018].

18. Sanders, J., Mayford, M., and Jeste, D. (2013). Empathic fear responses in mice are triggered by recognition of a shared experience. PLoS One 8, e74609. Available at: http://dx.plos.org/10.1371/journal.pone.0074609 [Accessed August 25, 2018].

19. Vogt, B. (2015). Chapter 21 - Cingulate Cortex and Pain Architecture. In The Rat Nervous System, G. Paxinos, ed. (London, U.K.: Academic PRess), pp. 575–596.

20. Allsop, S.A., Wichmann, R., Mills, F., Burgos-Robles, A., Chang, C.-J., Felix-Ortiz, A.C., Vienne, A., Beyeler, A., Izadmehr, E.M., Glober, G., et al. (2018). Corticoamygdala Transfer of Socially Derived Information Gates Observational Learning. Cell 173, 1329–1342.e18. Available at: http://linkinghub.elsevier.com/retrieve/pii/S0092867418304574 [Accessed June 8, 2018].

21. LeDoux, J.E. (2000). Emotion Circuits in the Brain. Annu. Rev. Neurosci. 23, 155–184. Available at: http://www.annualreviews.org/doi/10.1146/annurev.neuro.23.1.155 [Accessed July 30, 2018].

22. Supèr, H., and Roelfsema, P.R. (2005). Chronic multiunit recordings in behaving animals: advantages and limitations. Prog. Brain Res. 147, 263–82. Available at: http://linkinghub.elsevier.com/retrieve/pii/S0079612304470204 [Accessed July 2, 2018].

23. Zhang, Y., Wang, N., Wang, J.-Y., Chang, J.-Y., Woodward, D.J., and Luo, F. (2011). Ensemble encoding of nociceptive stimulus intensity in the rat medial and lateral pain systems. Mol. Pain 7, 64. Available at: http://journals.sagepub.com/doi/10.1186/1744-8069-7-64 [Accessed September 17, 2018].

24. Fan, R.-J., Kung, J.-C., Olausson, B.A., and Shyu, B.-C. (2009). Nocifensive behaviors components evoked by brief laser pulses are mediated by C fibers. Physiol. Behav. 98, 108– Available at: http://www.ncbi.nlm.nih.gov/pubmed/19410593 [Accessed September 17, 2018].

25. LeDoux, J.E. (2014). Coming to terms with fear. Proc. Natl. Acad. Sci. 111, 2871–2878. Available at: http://www.ncbi.nlm.nih.gov/pubmed/24501122 [Accessed August 1, 2018].

26. Cruccu, G., Sommer, C., Anand, P., Attal, N., Baron, R., Garcia-Larrea, L., Haanpaa, M., Jensen, T.S., Serra, J., and Treede, R.-D. (2010). EFNS guidelines on neuropathic pain assessment: revised 2009. Eur. J. Neurol. 17, 1010–1018. Available at: http://www.ncbi.nlm.nih.gov/pubmed/20298428 [Accessed September 5, 2018].

27. Bromm, B., and Treede, R.D. (1991). Laser-evoked cerebral potentials in the assessment of cutaneous pain sensitivity in normal subjects and patients. Rev. Neurol. (Paris). 147, 625–43. Available at: http://www.ncbi.nlm.nih.gov/pubmed/1763252 [Accessed September 5, 2018].

28. Jourdan, D., Ardid, D., Chapuy, E., Eschalier, A., and Le Bars, D. (1995). Audible and ultrasonic vocalization elicited by single electrical nociceptive stimuli to the tail in the rat. Pain 63, 237–49. Available at: http://www.ncbi.nlm.nih.gov/pubmed/8628590 [Accessed September 18, 2018].

29. Kohler, E., Keysers, C., Umiltà, M.A., Fogassi, L., Gallese, V., and Rizzolatti, G. (2002). Hearing sounds, understanding actions: action representation in mirror neurons. Science 297, 846–8. Available at: http://www.ncbi.nlm.nih.gov/pubmed/12161656 [Accessed November 14, 2015].

30. Gallese, V., Fadiga, L., Fogassi, L., and Rizzolatti, G. (1996). Action recognition in the premotor cortex. Brain 119 (Pt 2, 593–609. Available at: http://www.ncbi.nlm.nih.gov/pubmed/8800951.

31. Keysers, C., and Gazzola, V. (2017). A Plea for Cross-species Social Neuroscience. Curr. Top. Behav. Neurosci. 30, 179–191. Available at: http://www.ncbi.nlm.nih.gov/pubmed/26946502 [Accessed March 10, 2016].

32. Keysers, C., and Gazzola, V. (2009). Expanding the mirror: vicarious activity for actions, emotions, and sensations. Curr. Opin. Neurobiol. 19, 666–71. Available at: http://www.ncbi.nlm.nih.gov/pubmed/19880311 [Accessed December 5, 2015].

33. Paxinos, G., and Watson, C. (2007). The rat brain in stereotaxic coordinates 6th Editio. (Amsterdam, The Netherlands: Elsevier Academic Press).

34. Zhang, Q., Xiao, Z., Huang, C., Hu, S., Kulkarni, P., Martinez, E., Tong, A.P., Garg, A., Zhou, H., Chen, Z., et al. (2018). Local field potential decoding of the onset and intensity of acute pain in rats. Sci. Rep. 8, 8299. Available at: http://www.nature.com/articles/s41598-018-26527- w [Accessed September 17, 2018].

